# Pervasive *cis* effects of variation in copy number of large tandem repeats on local epigenetics and gene expression

**DOI:** 10.1101/2020.12.16.423078

**Authors:** Paras Garg, Alejandro Martin-Trujillo, Oscar L. Rodriguez, Scott J. Gies, Bharati Jadhav, Andrew J. Sharp

## Abstract

Variable Number Tandem Repeats (VNTRs) are composed of large tandemly repeated motifs, many of which are highly polymorphic in copy number. However, due to their large size and repetitive nature, they remain poorly studied. To investigate the regulatory potential of VNTRs, we used read-depth data from Illumina whole genome sequencing to perform association analysis between copy number of ~70,000 VNTRs (motif size ≥10bp) with both gene expression (404 samples in 48 tissues) and DNA methylation (235 samples in peripheral blood), identifying thousands of VNTRs that are associated with local gene expression (eVNTRs) and DNA methylation levels (mVNTRs). Using large-scale replication analysis in an independent cohort we validated 73-80% of signals observed in the two discovery cohorts, providing robust evidence to support that these represent genuine associations. Further, conditional analysis indicated that many eVNTRs and mVNTRs act as QTLs independently of other local variation. We also observed strong enrichments of eVNTRs and mVNTRs for regulatory features such as enhancers and promoters. Using the Human Genome Diversity Panel, we defined sets of VNTRs that show highly divergent copy numbers among human populations, show that these are enriched for regulatory effects on gene expression and epigenetics, and preferentially associate with genes that have been linked with human phenotypes through GWAS. Our study provides strong evidence supporting functional variation at thousands of VNTRs, and defines candidate sets of VNTRs, copy number variation of which potentially plays a role in numerous human phenotypes.

## INTRODUCTION

Tandem Repeats (TRs) are stretches of DNA comprised of two or more contiguous copies of a sequence of nucleotides arranged in head-to-tail pattern, e.g. CAG-CAG-CAG. The human genome contains >1 million TRs which collectively span ~3% of our total genome.^1^ These TRs range in motif size from mono-nucleotide repeats at one extreme (e.g. TTTTTTT), to those with much larger motifs that can in some cases be several kilobases in size, even containing entire exons or genes within each repeated unit.^2,3^ Due to their repetitive nature, TRs often show high mutation frequencies, with many showing extremely high levels of length polymorphism.^4,5^ For example, a recent comprehensive study of genome variation showed that ~50% of insertion-deletion events within the human genome map to TR regions. ^6^ However, despite contributing to a large fraction of genetic variation, TRs remain poorly studied, and as a result their influence on human phenotypes is almost certainly under-estimated. This is largely due to their repetitive and highly variable nature which, until the recent advent of specialized algorithms designed to genotype TR lengths from sequencing data, made them largely inaccessible to high throughput genotyping approaches.^7–12^

Previously, we and others have demonstrated functional effects on local gene expression and epigenetics of length variation in TRs with both short motifs (motif size 1-6bp, often termed microsatellites) and TRs with very large motifs (motif size >2kb, also termed macrosatellites).^13–16^ In contrast, TRs with motif sizes between these two extremes, often termed Variable Number Tandem Repeats (VNTRs) or minisatellites, have been less well studied. This is largely due to technical difficulties of genotyping variation at loci composed of moderate-to-large tandem repeats motifs, and is further compounded by the fact that many tandem repeats undergo a relatively high rate of recurrent mutation, meaning that copy number variation of large TRs is often poorly tagged by flanking SNVs.^16^ As a result, variation of many TR loci is poorly ascertained by standard SNV-based genome-wide association studies (GWAS). Thus, there is currently a knowledge gap regarding the role of TR variation in human disease.

Numerous targeted studies in the literature have implicated length variation of VNTR loci as putative drivers of human molecular and disease phenotypes. Specific examples include a 12mer repeat upstream of the *CSTB* gene [MIM: 601145] that is the strongest known expression quantitative trait locus (eQTL) associated with *CSTB* expression, a 30mer repeat in the promoter of *MAOA* [MIM: 309850] implicated in multiple neurologic and behavioral phenotypes, a 14mer repeat upstream of the *INS* gene [MIM: 176730] that is associated with multiple metabolic traits, insulin production and diabetes risk, and an 25mer repeat intronic within *ABCA7* [MIM: 605414] that is enriched for long alleles in Alzheimer disease and correlates with *ABCA7* splicing and amyloid β levels in cerebrospinal fluid.^17–22^

Building on this prior work, here we used read depth from Illumina whole genome sequencing (WGS) data to perform a genome wide analysis of copy number variation at ~70,000 VNTR loci (defined here as tandem repeats with motif size ≥10bp and span ≥100bp in the reference genome) in two discovery cohorts and a third replication population. Our study provides functional insight into a previously understudied fraction of human genetic variation, and suggests that future studies of VNTR variation may explain some of the ‘missing heritability’ of the human genome.^23,24^

## METHODS

### Description of cohorts used for VNTR association analysis

#### GTEx

We obtained Illumina 150bp paired-end WGS data and resulting variant calls made using GATK in 620 individuals from The Genotype-Tissue Expression (GTEx) project from dbGAP (Accession ID phs000424.v7.p2). RNAseq data for these samples were downloaded from GTEX portal (Version 7, https://www.gtexportal.org/), comprising quality-controlled and processed files for 48 tissues generated by the GTEx Consortium. These data were aligned to hg19, had already undergone filtering to remove genes with low expression, and been subject to rank based inverse normal transformation.

#### PCGC

WGS and methylation data for 249 individuals were selected from the cohort collected by the Pediatric Cardiac Genomic Consortium (PCGC). An extensive description of PCGC samples as well as further details about sample collection can be found in a summary publications released by the PCGC.^25,26^ Briefly, the cohort comprises individuals aged from newborn to 47 years (mean 8.2 years) diagnosed with a range of congenital heart defects, with conotruncal and left-sided obstructive lesions being the two most common diagnoses. Illumina 150bp paired-end WGS data generated using PCR-free libraries from peripheral blood DNA (average of 36x genome coverage, range 25-39x) were obtained from dbGAP (Accession ID phs001138.v1.p2). Peripheral blood methylomes were downloaded from GEO (Accession ID GSE159930), and normalized as described previously.^27^ We utilized the array data to infer the likely sex of each sample, based on scatter plots of mean β-value of probes located on chrX versus the fraction of probes located on chrY with detection p>0.01. We compared these predictions against self-reported gender for each sample, and removed four samples with a potential sex mismatch. We utilized data from autosomal probes, excluding any that mapped to multiple genomic locations. We also utilized the genotypes obtained from GATK analysis of the WGS data, and in each sample excluded β-values for any CpG that contained an SNV either within the probe binding site or the interrogated CpG. After these filters, a total of 821,035 CpG sites were retained for downstream analysis.

#### PPMI

We utilized data from the Parkinson’s Progression Markers Initiative (PPMI) cohort (https://www.ppmi-info.org/), corresponding to 712 individuals (189 healthy controls and 523 samples with varying types of Parkinsonism) with available Illumina WGS data aligned to the hg38 reference genome.^28^ RNAseq data generated from peripheral blood were available for 676 PPMI samples, comprising read counts for 22,582 genes listed in GENCODE v19. The read counts were filtered, normalized and subject to rank-based inverse normal transformation using scripts provided by the GTEx consortium (https://github.com/broadinstitute/gtex-pipeline). DNA methylation data generated using the Illumina 850k array from peripheral blood DNA were available for 524 PPMI samples.

### Estimation of VNTR copy number in two discovery cohorts

We downloaded 886,954 autosomal tandem repeats listed in the Simple Repeats track from the hg19 build of the UCSC genome browser (http://genome.ucsc.edu/), retaining only those repeats with motif size ≥10bp and total length of repeat tract ≥100bp. Where multiple tandem repeat annotations overlapped, these were merged together, resulting in 89,893 unique VNTR regions that were used in subsequent analysis. We did not include the sex chromosomes in our analysis.

In each sample of the discovery cohort (GTEx and PCGC), we estimated relative diploid copy number of each autosomal VNTR region using *CNVnator* (version 0.3.3, using default thresholds and bin size 100bp), which uses normalized read depth to estimate copy number of a locus.^29^ It should be noted that in VNTR regions, where by definition there are multiple copies of a repeated motif, *CNVnator* copy number estimates represent the fold change in total (diploid) repeat number relative to the number of motifs annotated in the (haploid) reference genome. For example, Figure 1A shows *CNVnator* estimated copy number for a 44mer repeat which has 43 copies in the reference genome (chr12:132,633,436-132,635,309). An individual with a relative *CNVnator* copy estimate of 6 is therefore predicted to carry a total of 43×6=258 copies of this repeat.

**Figure 1.**
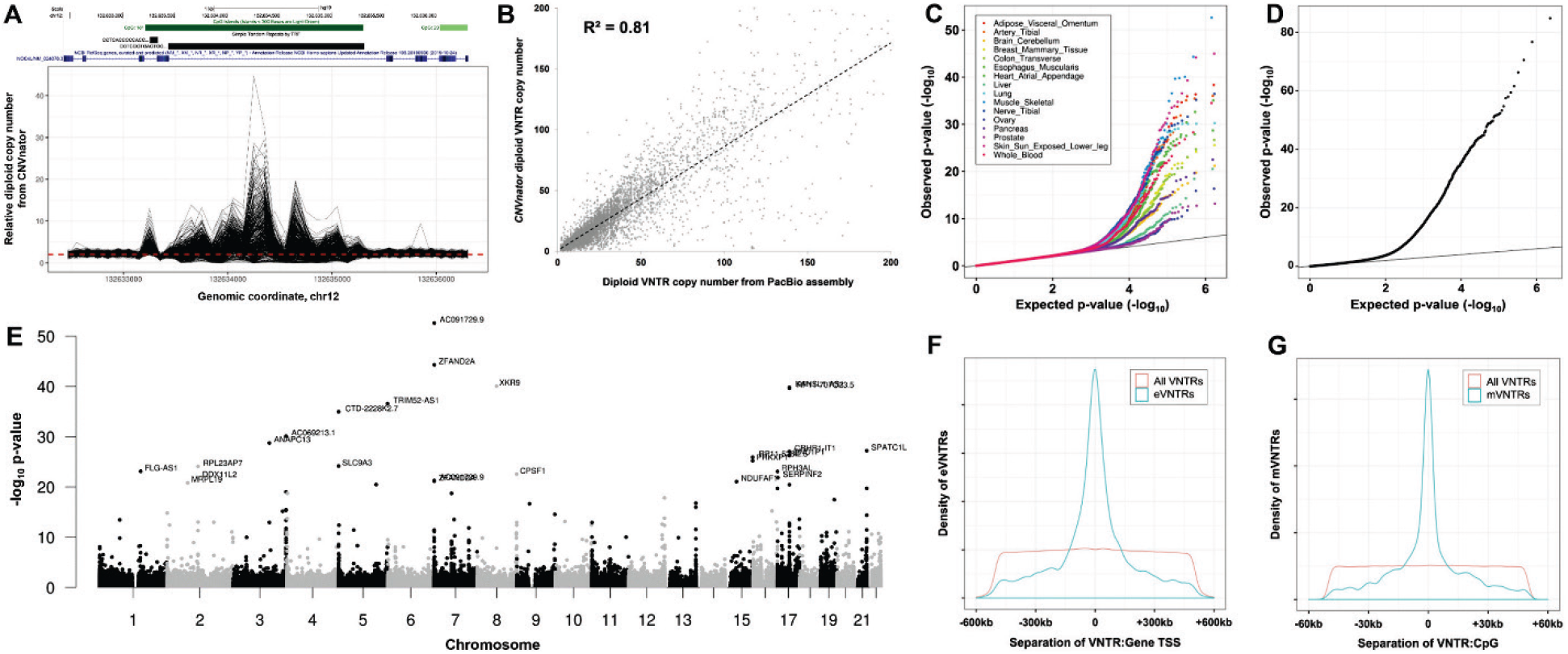
Copy number variation at thousands of VNTRs is associated with variation in gene expression and DNA methylation *in cis*. **(A)** *CNVnator* estimated copy number per 100bp bin over a VNTR region shows highly variable read depth among samples from the GTEx cohort. Shown is read depth data for a 44mer repeat which has 43 copies in the reference genome (chr12:132,148,891-132,150,764, hg38), located intronic within the *NOC4L* gene, which shows >10-fold difference in copy number within the population. **(B)** Read depth provides good accuracy for estimating diploid VNTR copy number. Using 14 samples where both Illumina and PacBio WGS data were available, at 1,892 eVNTR loci we compared diploid VNTR copy number estimates from WGS read depth using *CNVnator* with direct genotypes derived from Pacific Biosciences long read diploid assemblies. We observed a high correlation between the two approaches (R^2^=0.81). **(C)** QQ plots showing the distribution of observed versus expected p-values for eVNTRs in 16 representative GTEx tissues, and **(D)** mVNTRs in whole blood from the PCGC. Variations in the observed p-value distribution among GTEx tissues are a reflection of the varying sample sizes available, which strongly influences statistical power. **(E)** Manhattan plot showing results of *cis*-association analysis between VNTR copy number and gene expression in skeletal muscle samples from the GTEx cohort. The high frequency of significant associations in subtelomeric and centromeric regions is consistent with the known enrichment of VNTRs in these regions.^59,60^ **(F,G)** Significant eVNTRs and mVNTRs are highly enriched within close proximity to the genes and CpGs whose expression or methylation level they are associated with, respectively, mirroring similar observations made for SNV eQTLs and mQTLs.^61,62^ However, we note an approximate order of magnitude difference in the distances over which significant eVNTRs and mVNTRs were typically observed to function.

Utilizing *CNVnator* copy number estimates of invariant regions of the genome, we observed a strong technical bias in GTEx WGS data, with samples that were sequenced prior to 2016 showing systematic shifts in estimated copy number compared to later batches (Figure S1). As a result, we removed 136 samples that were sequenced in batch “2015-10-06”. Based on analysis of invariant loci, and principal component analysis and density plots based on VNTR copy number, we excluded a further 60 samples from the GTEx and 10 samples from the PCGC cohorts, respectively, that were outliers in one or more of these analyses (Figure S2).

In situations where a VNTR is embedded within a larger copy number variable region, copy number estimates for a VNTR based on read depth can be confounded by variations of the wider region, as these would result in gains or losses in the total number of VNTR copies present, but without any change in the length of the VNTR array. To identify VNTRs where our copy number estimates were potentially subject to this confounder, we performed copy number analysis of the 3’ and 5’ regions flanking each VNTR using *CNVnator* (Figure S3). In cases where the 1kb flanking region of a VNTR overlapped a simple repeat with motif size ≥6bp, we trimmed the flanking region, retaining only the flanking portion that was adjacent to the VNTR. We then removed from our analysis any VNTRs where:

1. Both flanks had trimmed length <500bp
2. Correlation (R) between copy number of the VNTR and either of the flanking regions was >0.5
3. Either flanking region showed large variations in copy number, defined as those flanks where the difference between 99^th^ and 1^st^ percentile was >2
4. We also removed any VNTRs that overlapped CNVs with minor allele frequency >10% in Europeans.^30^

As copy number estimates in GTEx WGS samples showed high variability based on analysis of density plots, we normalized VNTR copy numbers in the 424 remaining samples by applying an inverse rank normal transformation.^31^ Based on visual inspection of density plots of these transformed copy numbers, we removed a further 20 outlier samples (Figure S2), leaving 404 samples that were used for association analysis with gene expression (Table S1). Finally, we removed VNTRs that showed very low levels of variation in the population (standard deviation <0.2).

### Comparison of VNTR copy number estimates with genotypes obtained using long reads and the adVNTR algorithm

Using the tool MsPAC, we generated diploid genome assemblies for 14 individuals from available Pacific Biosciences (PacBio) WGS data and phased SNVs (Table S2).^32^ Where phased SNVs were not available (samples HG02059, HG02818, HG03486 and HG0386), we performed phasing using *GATK HaplotypeCaller* and *WhatsHap*.^33^ We generated VNTR genotypes directly from the diploid long-read assemblies using *PacMonSTR*.^34^ For each of these individuals, PCR-free Illumina WGS data were also available, and were processed with *CNVnator* to estimate VNTR copy number, as described above. To estimate the accuracy of our VNTR genotypes derived using *CNVnator*, we utilized a set of 2,027 eVNTRs that showed significant associations with gene expression in one or more GTEx tissues, and which were composed of single annotated (i.e. non-merged) tandem repeats, copy number of which could be unambiguously genotyped using *PacMonSTR*. We discarded genotypes where both haplotypes in a sample were not represented in the PacBio genome assemblies, or where VNTR copy number was >200, yielding a final total of 16,403 pairwise genotypes derived from 1,891 VNTR loci across the 14 samples, representing all eVNTR loci genotyped by *CNVnator* for which we also obtained at least one set of diploid genotypes from the 14 PacBio genome assemblies analyzed. To assess the performance of an alternative approach for genotyping VNTRs from short read WGS, we were also able to generate genotypes for 1,746 of these same 1,891 loci from the Illumina WGS reads with *adVNTR* in the 14 samples using default parameters.^11^

### Identification of eVNTRs in the GTEx cohort

Using RNAseq data from the filtered set of 404 WGS samples that passed our quality control steps, gene expression data were adjusted for gender, RNAseq platform, the first three principal components from SNV genotypes, and between 15-60 covariates per tissue estimated using PEER.^35^ Within each tissue, we performed linear regression between VNTR copy number and corrected expression level of each gene located within ±500kb using the lm function in R. We applied a False Discovery Rate correction, and reported all VNTR:gene pairs with FDR q<0.1 in any tissue.^36^

### Identification of mVNTRs in the PCGC cohort

After excluding samples that either did not pass our QC for DNA methylation or were outliers for VNTR copy number based on density plots, 235 individuals from the PCGC cohort were utilized for association analysis of VNTR copy number with CpG methylation levels. We excluded CpGs with low levels of variation (standard deviation <0.02), leaving 316,169 CpGs that were located within ±50kb of VNTRs that were used for association analysis. CpG methylation data (β-values) were adjusted for age, sex, the top two ancestry-related principle components derived from PCA of SNVs, and blood cell fractions estimated directly from the methylation data using the Houseman method.^37,38^ The resulting residuals were used to test the association between DNA methylation and estimated VNTR copy number using the lm function in R. We applied a Bonferroni correction to the resulting p-values based on the total number of pairwise VNTR:CpG tests performed, and considered those with Bonferroni adjusted p<0.01 as significant.

### Replication of eVNTRs and mVNTRs in the PPMI cohort

We utilized available WGS, RNAseq and methylation data for 712 individuals from the PPMI cohort. We generated copy number estimates for all VNTR loci utilized in the GTEx and PCGC discovery cohorts using *CNVnator* (version 0.4.1), and applied the same quality control and normalization steps as used in the discovery cohorts, including inverse rank normal transformation to the VNTR copy numbers, resulting in the exclusion of nine outlier samples.

We normalized gene expression data using the same method as applied to the GTEx cohort, including application of inverse rank normal transformation. These normalized expression data were adjusted for gender, the first three ancestry-related principle components derived from PCA of SNVs, and 60 additional components estimated using PEER.^35^ We performed association between VNTRs and normalized adjusted gene expression levels using linear regression, as described above for the GTEx cohort.

For replication of mVNTRs, we applied the same quality control and normalization pipeline to the methylation data as used for the PCGC cohort, as described above. Normalized β-values were adjusted for gender, age, the top three principal components from SNV genotypes, and estimated blood cell fractions. The residuals were then used to perform linear regression with VNTR genotype.

### Enrichment analysis

All enrichment analyses were performed by comparing the frequency of significant eVNTRs and mVNTRs against the background set of all VNTR:gene pairs that were tested in each cohort, with p-values generated using the hypergeometric distribution. We defined promoter regions as ±2kb of gene transcription start sites (TSS). We utilized a set of enhancer elements downloaded the GeneHancer track in the UCSC Genome Browser, utilizing only loci labelled “Enhancers”.^39^ We utilized a composite list of silencer elements corresponding to all significant silencer elements identified in two cell types under different conditions.^40^

### Population stratification of VNTRs

We obtained Illumina WGS reads mapped to hg38 from samples in the Human Genome Diversity Panel, utilizing data for a subset of 676 samples that were sequenced using PCR-free protocols.^41^ We used *CNVnator* (version 0.4.1) to estimate relative copy number of autosomal VNTRs (hg38). We performed quality control using PCA and density plots to remove outliers, and compared the reported gender of each sample against sex chromosome copy number, removing any discordant samples. We filtered VNTRs to remove those within putative larger CNVs, as detailed above. After applying these filters, we utilized genotypes of 66,796 VNTRs in 643 samples from seven different super-populations. For each super-population, we calculated V_ST_ as follows:

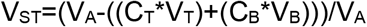

Where V_A_ is the variance of all the samples, V_T_ is the variance of the target population, V_B_ is the variance of the background population, while C_T_ and C_B_ are fractions of the number of target and background population, respectively.^42^ For each of the seven super-populations, we calculated V_ST_ for each VNTR by considering one super population as the target, and using all other samples as background. P-values were generated by permutation testing (n=1,000 permutations), with samples randomly assigned to either the target or background groups. We selected those VNTRs in each super-population with V_ST_≥0.1 and permutation p<0.01.

### Annotation of VNTRs with potential trait associations

In order to link eVNTRs with human traits that they might influence, we used two complementary approaches. First, we used results of *PrediXcan* applied to 44 GTEx tissues and >100 phenotypes from GWAS, annotating eVNTRs with phenotypes if they shared the same gene name and tissue as indicated by *PrediXcan*.^43^ However, as *PrediXcan* has been applied to a relatively limited set of traits, we further annotated eVNTRs using a combination of eQTLs identified by the GTEx project and SNVs from the GWAS Catalog.^44^ Here, eVNTRs were linked to putative associated phenotypes as follows: (i) For each eVNTR identified in a specific tissue, we joined these with eQTLs identified in the same GTEx tissue based on gene name; (ii) We extracted all SNVs from the GWAS Catalog with p<5×10^−8^, and joined these to the GTEx eQTLs (iii) Where an eVNTR was joined with an SNV that was both a GWAS variant and an eQTL for the same gene in the same tissue, we annotated the eVNTR with the GWAS phenotype(s).

## RESULTS

### Robust genotyping of VNTRs using read depth

Using read depth from Illumina WGS data as a proxy for diploid copy number, we generated copy number estimates for a set of 70,787 large tandem repeats (median motif size 116bp, mean span of repeat tract in reference genome 353bp), henceforth referred to as Variable Number Tandem Repeats (VNTRs). Many VNTR loci showed highly variable copy number estimates among different individuals, indicative of extreme levels of inter-individual polymorphism at many of these loci (Figure 1A).

In order to assess the validity of genotyping VNTRs from read depth, we compared estimated VNTR copy numbers from *CNVnator* with genotypes obtained directly from spanning long reads from *de novo* diploid PacBio genome assemblies. Using 14 individuals for which both Illumina and PacBio WGS data were available, we observed good global correlation between these two approaches, with an overall R^2^=0.81, indicating that read depth is generally an effective proxy for measuring total copy number at the majority of VNTR loci (Figure 1B, Table S3). In comparison, we found that an alternative tool designed for genotyping VNTRs from short read data performed relatively poorly, yielding an R^2^=0.14 when compared with direct genotypes generated from long-read WGS (Figure S4, Table S3).^11^

Given that some VNTR motifs are not unique and can occur at multiple genomic loci, we investigated the reliability of reads mapped to VNTR loci. Using high coverage Illumina WGS data in a Yoruban individual from the 1000 Genomes Project (NA18874), we assessed mapping quality scores for reads that overlapped VNTRs based on both their MAPQ score, and the MAPQ score of their mate-pairs. We classified reads from VNTR loci into three categories: (i) MAPQ≥10, which we considered reliably mapped, (ii) MAPQ<10, but with a mate-pair that mapped reliably within ±10kb. We considered these reads as reliably mapped to the correct VNTR on the basis of their mate pair. Likely many such reads that are contained entirely within a VNTR yield low mapping quality due to the fact that VNTRs are composed of repeated copies, giving multiple possible map positions within a single VNTR tract. (iii) MAPQ<10, and with a mate pair that was not anchored within ±10kb. We considered these reads unreliably mapped. Overall, we observed that the vast majority of reads from VNTR loci were reliably mapped: 97.5% of VNTRs comprised <10% of overlapping reads that were unreliably mapped (MAPQ<10 and no anchoring mate pair), and only a single VNTR contained >50% of unreliably mapped reads (Figure S5). These data indicate that ambiguous read mapping to tandemly repeated regions is not a significant confounder of our approach.

### Overview of association analysis of VNTRs with gene expression and DNA methylation

To assess the potential regulatory effects of copy number changes of VNTRs on local gene expression and epigenetics we utilized two discovery cohorts for which PCR-free Illumina WGS data were available: (i) a subset of quality-filtered samples from the GTEx project, comprising 404 individuals with expression data from 48 different tissues. Here we performed *cis*-association analysis between estimated VNTR copy number and normalized gene expression within ±500kb. (ii) 235 quality-filtered samples from the PCGC for which DNA methylation profiles from whole blood were available. Here we performed *cis*-association analysis between estimated VNTR copy number and CpG methylation levels within ±50kb.

### Summary of gene expression associations in the GTEx cohort

After multiple testing correction, in the GTEx cohort we identified a total of 13,752 significant pairwise VNTR:gene expression associations (10% FDR) across 48 different tissues, corresponding to 2,980 unique expression QTL VNTRs (henceforth termed eVNTRs) that were associated with the expression level of 3,167 different genes (Table S4). Using QQ-plots to explore the distribution of observed versus expected associations, in each GTEx tissue we observed a clear enrichment for significant associations compared to the null distribution, with little evidence of genomic inflation (mean λ=1.019, range 0.997-1.040) (Figure 1C). As expected, the number of significant associations observed in different tissues was strongly associated with sample size (i.e. statistical power), varying from 13 identified in uterus to 1,080 in thyroid (Table S4). An example of the distribution of genome-wide association signals observed in skeletal muscle is shown in Figure 1E. Of note, we frequently observed the same VNTR:gene pairwise associations in multiple different tissues (35% of VNTR:gene associations were seen in ≥2 tissues), and of these, 99.4% showed consistent directionality in different tissues (Figure S6, Table S4). In addition, 34% of eVNTRs were associated with the expression of multiple different genes (mean of 3 associated genes per eVNTR, range 1-48).

Supporting a biological role in modulating gene expression, eVNTRs showed enrichments for several genome annotations with regulatory potential. We observed a 7.9-fold enrichment for eVNTRs located within ±2kb of transcription start sites (p=1.1×10^−73^, Figure 1F). Consistent with this observation, the sequence content of eVNTRs also showed a strong bias towards higher GC-content (permutation p<10^−7^) (Figure S7). We also observed that eVNTRs were enriched at both annotated enhancers (1.8-fold enrichment, p=2.0×10^−31^) and silencer elements (2.5-fold enrichment, p=1.9×10^−4^). Further examples of results observed at eVNTRs in the GTEx cohort are shown in Figure S8.

In further support of our results, we successfully replicated three associations of VNTRs with the expression level of individual genes that had been identified in previous targeted studies: a 36mer coding VNTR in exon 1 of *AS3MT* [MIM: 611806] that is associated with *AS3MT* expression and schizophrenia risk, a 72mer intronic VNTR that regulates *SIRT3* expression [MIM: 604481], and a 33mer promoter VNTR that regulates expression of *TRIB3* [MIM: 607898], a gene which has been linked with multiple human phenotypes.^45–47^

### Summary of DNA methylation associations in the PCGC cohort

After multiple testing correction, in the PCGC cohort we identified a total of 3,152 VNTR:CpG pairwise associations (Bonferroni-corrected p<0.01), corresponding to 1,480 unique methylation QTL VNTRs (henceforth termed mVNTRs) and 2,466 unique CpGs (Table S5). Similar to observations made for eVNTRs, mVNTRs also showed a strong bias to occur in close proximity to the CpGs they associated with, with the majority being separated by <5kb (Figure 1G). mVNTRs tended to have a significantly higher GC-content than all VNTRs in the genome (permutation p<10^−7^, Figure S7), were 2.4 fold enriched for enhancers (p=5.9×10^−34^), and 2.2-fold enriched for silencers (p=8.1×10^−3^). Three examples of the association signals observed around mVNTRs are shown in Figure 2, while additional plots of eight other mVNTR loci are shown in Figure S9.

**Figure 2.**
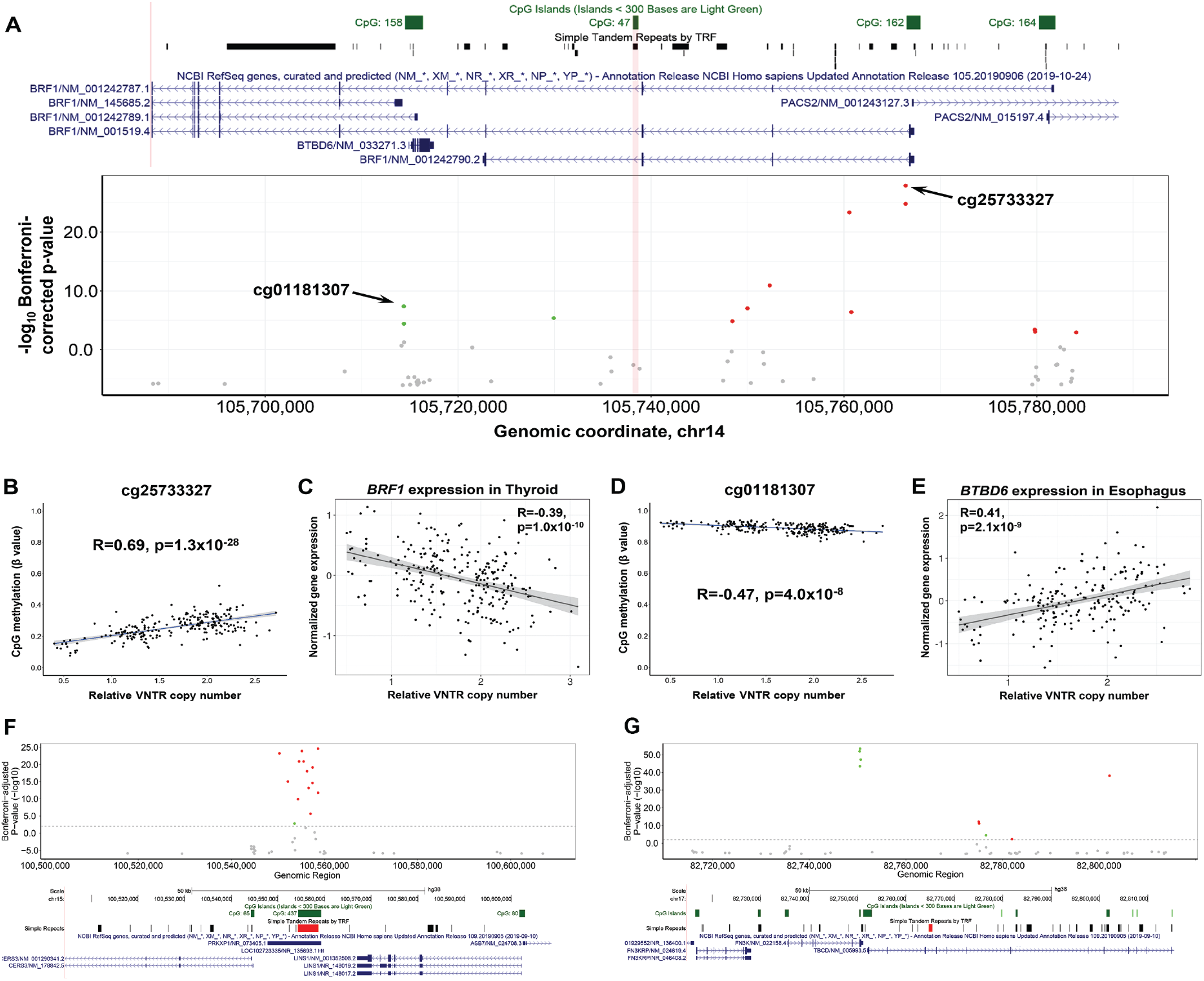
Example associations of VNTRs with *cis*-linked DNA methylation and gene expression. Copy number of a 72mer tandem repeat (chr14:105,271,805-105,272,305, hg38) is associated with DNA methylation levels at multiple CpGs spread over >80kb and the expression of multiple genes *in cis*. **(A)** Manhattan plot of associations between copy number of this VNTR and CpG methylation within ±50kb. Significant CpGs (p<0.01 after Bonferroni correction for the number of pairwise tests performed genome-wide), are shown in color, with red representing positive correlations with VNTR copy number, and green indicating negative correlations. The location of the 72mer VNTR is indicated by the vertical red bar in the center of the plot. Above the plot is an image from the UCSC Genome Browser showing location of CpG islands, Simple Repeats, and Refseq genes. **(B,C)** Correlation of VNTR copy number with CpG methylation (cg25733327), that lies 1kb downstream of the TSS of *BRF1,* and expression of *BRF1* in thyroid. **(D,E)** Correlation of VNTR copy number with CpG methylation (cg01181307) that lies 500bp upstream of the TSS of *BTBD6,* and expression of *BTBD6* in esophagus muscularis. For both genes, increased methylation levels around the TSS are associated with reduced gene expression, which is consistent with the known repressive effects of promoter methylation. **(F)** A 107mer repeat (chr15:100,554,293-100,558,659, hg38) increased copy number of which causes local hypermethylation. This VNTR also associates with the expression level of multiple nearby genes in many different tissues. **(G)** A 40mer repeat (chr17:82,764,738-82,765,449, hg38), which associates with methylation of multiple CpGs over an ~50kb region. This VNTR also associates with the expression level of multiple nearby genes in many different tissues. In F and G, the location of the associated VNTR is shown by a red bar.

### Conditional analysis indicates many VNTR associations are independent of SNV QTLs

Given that multiple different genetic variants may exert regulatory effects on gene expression and CpG methylation, we considered the possibility that the VNTR associations we observed might be indirect correlations driven by linkage disequilibrium between VNTRs and other variants that are the primary QTLs. To assess whether VNTRs act as QTLs independent of other local genetic variation, we performed conditional analyses by removing the effect of the strongest SNV QTL associated with each gene and CpG that were putatively associated with VNTR copy number.

First, we utilized SNV genotypes from the WGS data in our two discovery cohorts to identify SNVs that were significantly associated (FDR q<0.1) with local gene expression and CpG methylation levels (Figure 3A). For each VNTR pairwise association, we then retained only the subset of individuals that were homozygous for the major allele of the lead QTL SNV, and repeated the association analysis between VNTR copy number and gene expression/DNA methylation (Figure 3B). Doing so, we observed a clear trend where the majority of VNTR associations retained the same directionality as in our original analyses (Figures 3C and 3D). Overall, 9,791 of 12,784 eVNTR:gene pairs (76.6%), and 2,280 of 3,152 mVNTR:CpG pairs (72%) showed the same direction of association after conditioning on the lead QTL SNV. Despite a considerable loss of statistical power due to the reduced sample size when conditioning based on the strongest SNV QTL, in the GTEx cohort, 2,146 associations showed the same direction of effect with p<0.01, and 1,434 met our genome-wide significance discovery threshold (FDR q<0.1) (Table S4). Similarly, for mVNTRs identified in the PCGC cohort, after conditioning on the lead mQTL SNV, 693 associations showed the same direction of effect with p<0.01, and 273 associations met our genome-wide significance discovery threshold (Bonferroni p<0.01) (Table S5).

**Figure 3.**
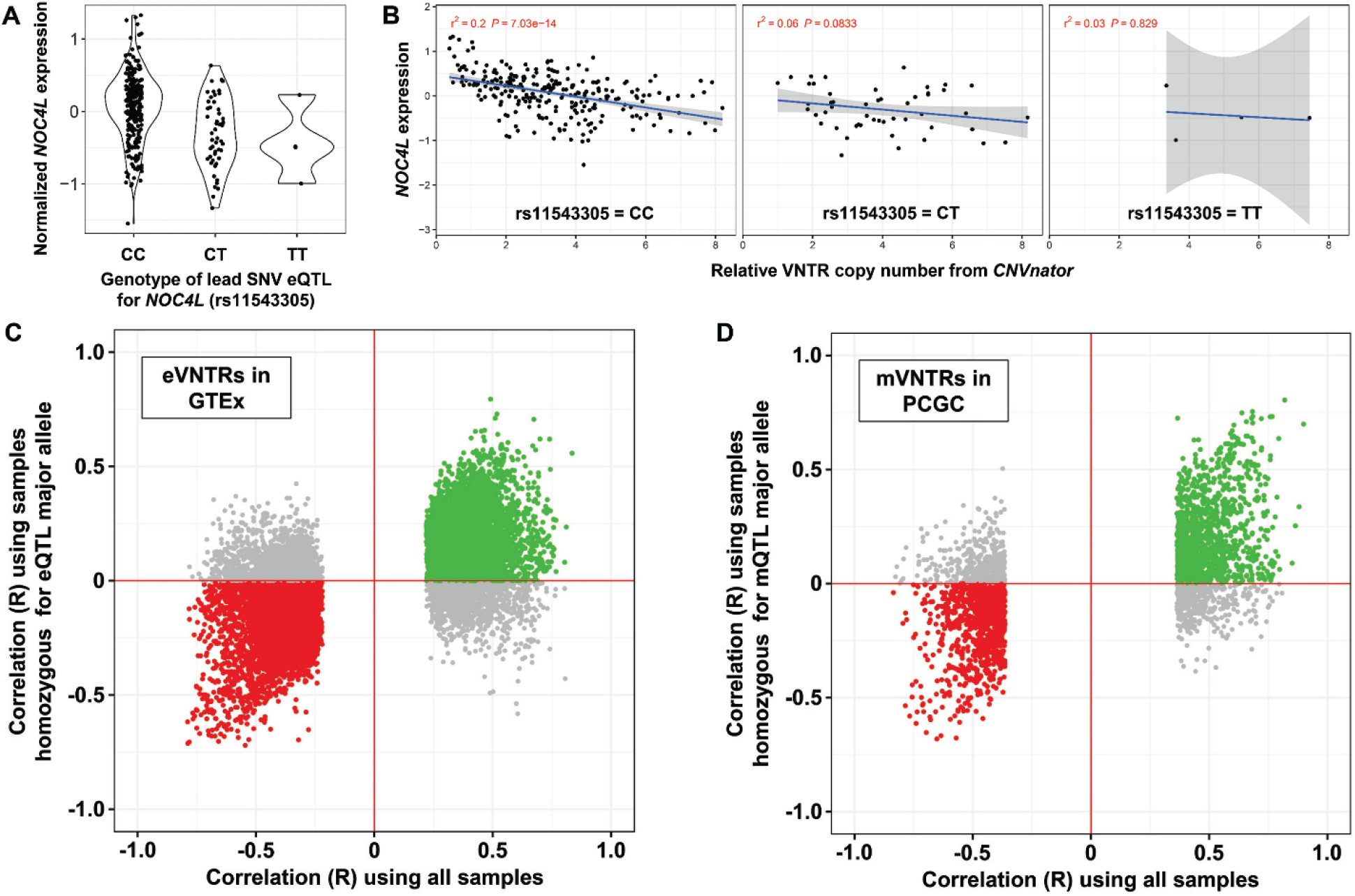
Copy number variation at the majority of VNTRs shows association with gene expression and DNA methylation independently of SNV eQTLs and mQTLs. In order to assess whether the observed associations we detected with VNTRs might simply be a result of linkage disequilibrium of VNTRs with local SNV haplotypes, we assessed the correlation of eVNTRs and mVNTRs with their target gene/CpG after conditioning on the strongest SNV which acts as a QTL on the same target. For each eVNTR:gene and mVNTR:CpG pair, we identified the peak SNV QTL for that gene or CpG, and then repeated the association test with VNTR copy number utilizing only those individuals that were homozygous for the major allele of the SNV QTL. Shown is an example locus, a 44mer repeat which has 43 copies in the reference genome (chr12:132,148,891-132,150,764, hg38), corresponding to the same VNTR shown in Figure 1A. This VNTR is located intronic within the *NOC4L* gene, and is significantly associated with *NOC4L* expression. **(A)** We identified rs11543305, a C/T variant which is located 1.6kb proximal to the VNTR, as being the lead SNV associated with *NOC4L* expression. **(B)** After stratifying samples based on genotype at rs11543305, copy number of this VNTR still shows a significant association with *NOC4L* expression. Considering all significant VNTRs we identified, including **(C)** eVNTRs observed in GTEx, and **(D)** mVNTRs observed in the PCGC cohort, there is a clear trend where the majority of observed VNTR associations retain their original signal even after are conditioning on the genotype of the lead SNV QTL. These data indicate that the majority of VNTR associations we identified act independently of local SNV QTLs. In each plot, colored points represent VNTR associations that retain the same directionality after conditioning on the lead SNV QTL, either positive associations (green) or negative associations (red).

Overall, these results indicate that many of the VNTR associations we detected are independent of other local QTLs, and are not simply driven by the linkage disequilibrium architecture of the genome.

### Large-scale replication of eVNTRs and mVNTRs in an independent cohort

In order to assess the robustness of the associations we identified in the GTEx and PCGC discovery cohorts, we conducted replication analysis in the PPMI cohort, consisting of a total of 703 individuals with WGS, gene expression and methylation data. We used *CNVnator* to analyze VNTR copy number in each sample, and then performed association analysis with both gene expression and CpG methylation levels using identical pipelines as applied in the two discovery cohorts. These analyses identified 3,537 significant eVNTRs that were associated with the expression level of 3,615 unique genes (6,454 pairwise associations) (Table S6), and 3,288 significant mVNTRs that were associated with methylation levels of 6,999 unique CpGs (9,730 pairwise associations) (Table S7).

When compared to the associations identified in whole blood from the GTEx and PCGC cohorts, we observed replication at genome-wide significance levels and with concordant directionality for 278 of 381 (73%) GTEx eVNTR:gene pairwise associations, and for 2,507 of 3,139 (80%) PCGC mVNTR:CpG pairwise associations (Figure 4), yielding strong evidence to support that the majority of associations we report are likely robust. We also observed a trend for many VNTR loci to be associated with both gene expression and CpG methylation. In the PPMI cohort, of the 3,537 unique eVNTR loci identified, 1,489 (42.1%) were also associated with the methylation level of one or more *cis*-linked CpGs. Of these, 653 (43.9%) had one or more associated CpGs that were located in either the promoter or an annotated enhancer element of the same gene whose expression they associated with. This high degree of convergence between these two data types lends further support to our results, and suggests that in at least a subset of cases, the potential mechanism of action of VNTRs on gene expression is via epigenetic modification of regulatory elements.

**Figure 4.**
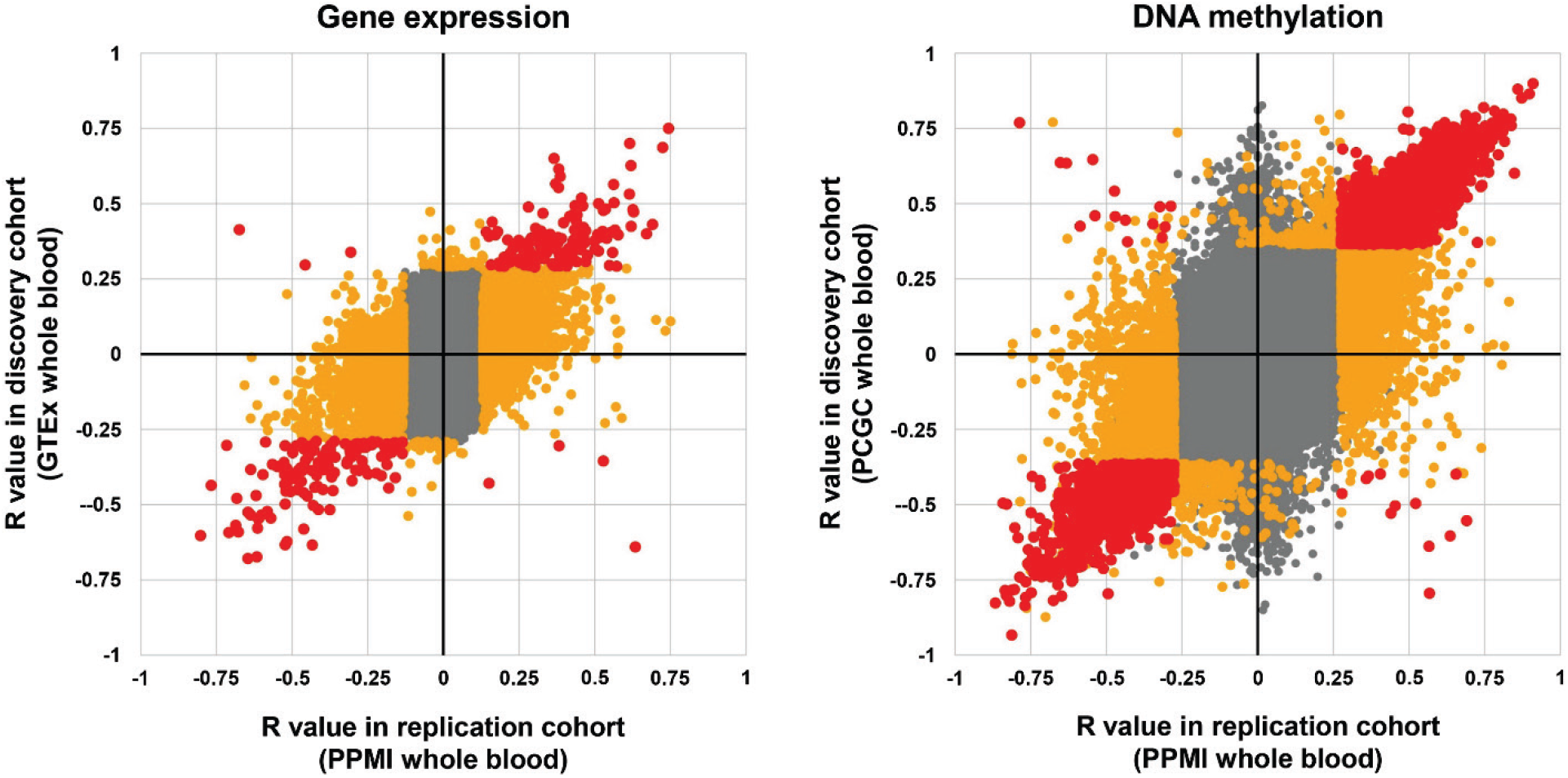
Replication of the majority of significant eVNTRs and mVNTRs in an independent cohort. In order to assess the robustness of the associations of VNTR copy number with gene expression and DNA methylation detected in our two discovery cohorts, we performed replication analysis in the PPMI cohort, which comprises 712 individuals with Illumina WGS, DNA methylation and RNAseq data derived from whole blood. We observed that 73% of significant eVNTRs detected in GTEx whole blood were also identified as significant in the PPMI cohort. Similarly, 80% of significant mVNTRs detected in the PCGC discovery cohort were also significant in the PPMI cohort.

### Population stratification and trait associations of VNTRs

We analyzed VNTR copy number in samples from the Human Genome Diversity Panel, using these data to estimate the degree of population stratification in VNTR copy number with the VST statistic.^41,42^ Examples of VNTRs with high population stratification are shown in Figure 5. We observed strong enrichment for VNTRs with high population divergence within the set of putatively functional VNTRs identified in our discovery cohorts: there were 27 GTEx eVNTRs with V_ST_>0.2 (5.7-fold enrichment compared to all VNTR loci tested, p=7.9×10^−14^), and 120 with V_ST_>0.1 (3.8-fold enrichment, p=9.2×10^−38^), while for mVNTRs in the PCGC cohort, 15 had V_ST_>0.2 (6.3-fold enrichment, p=1.3×10^−8^), and 112 had V_ST_>0.1 (6.6-fold enrichment, p=1.5×10^− 57^). We also compared this set of population-stratified VNTRs to tandem repeats that were previously identified as having expanded specifically in the human lineage compared to other primates and observed similar enrichments (GTEx eVNTRs 5.7-fold enriched, p=0.045; PCGC mVNTRs 9.2-fold enriched, p=0.018).^48^

**Figure 5.**
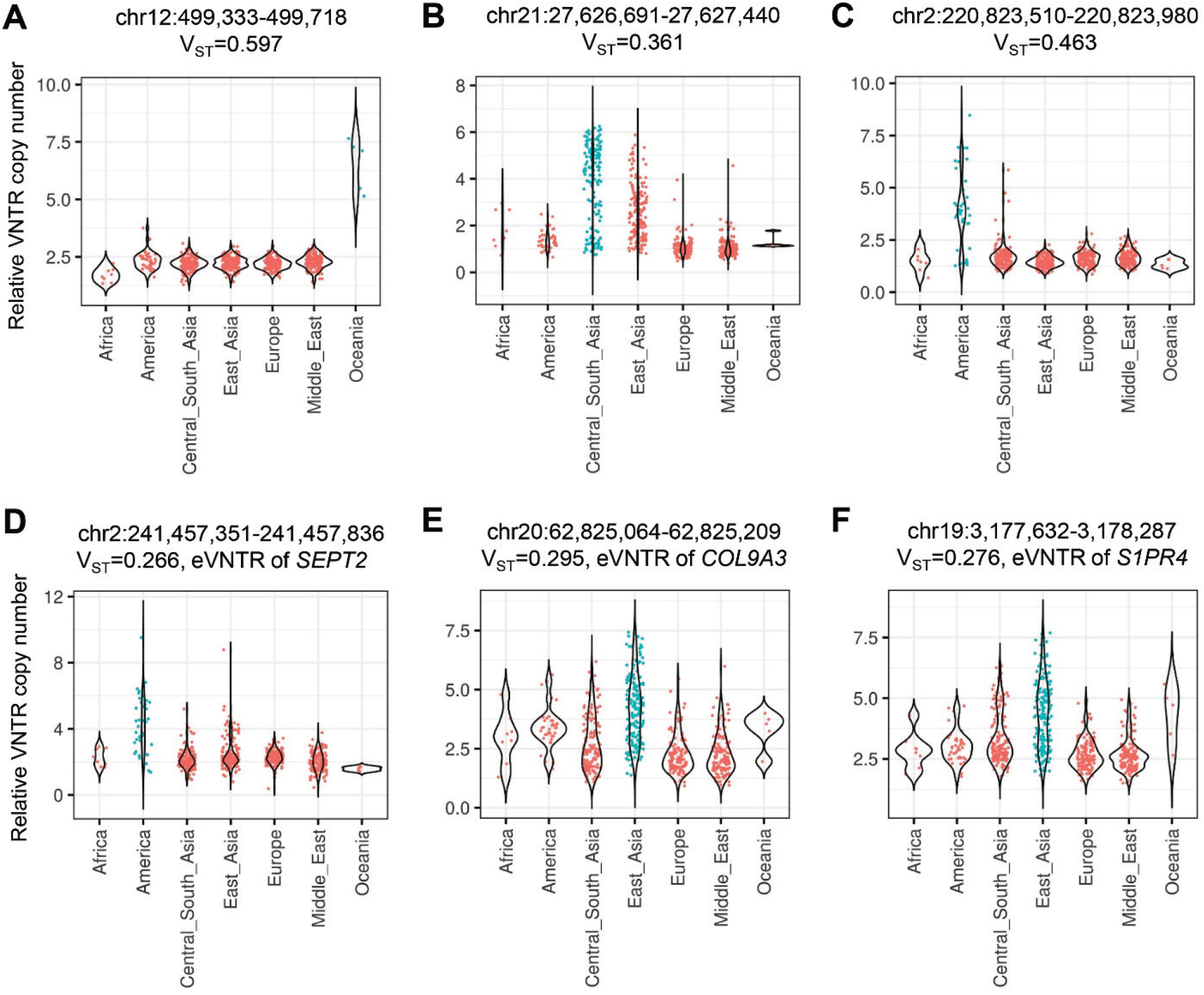
VNTRs with high population divergence are enriched for functional associations with gene expression, methylation, and human traits. We estimated population stratification of VNTR copy number with the V_ST_ statistic in samples from the Human Genome Diversity Panel. Both eVNTRs and mVNTRs were enriched for VNTRs with high V_ST_, and consistent with the notion that selection may have acted to modify copy number of functional VNTR loci in specific populations, we also observed that eVNTRs with elevated V_ST_ were enriched for putative phenotype associations. **(A-F)** Shown are six example VNTRs with high V_ST_. **(D)** An 81mer VNTR (chr2:241,457,351-241,457,836, hg38) showed expansion in Americans, is associated with expression level of *SEPT2* [MIM: 601506] in skin and thyroid, and is potentially linked to multiple human traits by GWAS studies. **(E)** A 24mer VNTR (chr20:62,825,064-62,825,209, hg38) showed expansion in East Asians, and is associated with expression level of *COL9A3* [MIM: 120270] in adipose tissue, muscle and blood. **(F)** A 39mer VNTR (chr19:3,177,632-3,178,287, hg38) showed expansion in East Asians, and is associated with expression level of *S1PR4* [MIM: 603751] in mammary tissue, thyroid and esophagus.

In order to investigate whether eVNTRs with elevated V_ST_ levels were enriched for phenotype associations, we annotated eVNTRs with human phenotypes that they potentially regulate using both the results of *PrediXcan*, and a combination of tissue-matched eQTLs joined with variants from the GWAS Catalog (Table S4). This identified 198 of 2,980 eVNTRs (6.6%) that had trait annotations from *PrediXcan*, while 634 eVNTRs (21.3%) had annotations derived from the overlap of GWAS Catalog variants and eQTLs. Examples of several functionally interesting candidate eVNTRs that are potentially linked to human traits using annotations from *PrediXcan* include:

i. An 87mer VNTR (chr6:166,997,608-166,997,912, hg38) which associates with expression of *RNASET2* [MIM: 612944] in esophagus mucosa. *RNASET2* is a secreted extracellular ribonuclease with roles in immune sensing and response, and is linked by *PrediXcan* with risk of Crohn’s, inflammatory bowel disease, and rheumatoid arthritis.^49,50^
ii. A VNTR region composed of multiple motifs (chr16:29196863-29197354, hg38) which associates with expression of *TUFM* [MIM: 602389] in thyroid, *TUFM* is a mitochondrial elongation factor involved in mitochondrial replication, and is linked by *PrediXcan* with body mass index, hip and waist circumference.^51^
iii. A 53mer VNTR (chr17:83,032,018-83,032,543, hg38) located intronic within *B3GNTL1* [MIM: 615337], which associates with *B3GNTL1* expression in aorta, *B3GNTL1* is a glycosyltransferase that transfers sugar moieties to acceptor molecules, and is linked by *PrediXcan* with levels of glycated hemogoblin.^52^

Consistent with the notion that selection may have acted to modify copy number of functional VNTR loci in specific populations, we observed that eVNTRs with elevated V_ST_ levels were enriched for putative phenotype associations: 44 GTEx eVNTRs with V_ST_>0.1 were linked with GWAS traits, representing a 1.7-fold enrichment when compared to all eVNTRs identified (p=9.0×10^−5^), while 13 had trait associations from *PrediXcan* (1.6-fold enrichment, p=0.058).

## DISCUSSION

Here we have conducted a genome-wide scan for putatively functional VNTRs that associate with local gene expression (eVNTRs) and DNA methylation (mVNTRs) using two separate cohorts for initial discovery, followed by subsequent replication in a third cohort. We identified thousands of VNTRs where repeat copy number associated with local expression and epigenetics, and successfully replicated the majority of these signals at stringent genome-wide significance thresholds. Multiple observations are consistent with a functional role for these loci, including an enrichment for regulatory elements such as gene promoters, enhancer and silencer elements, a strong bias for eVNTRs/mVNTRs to lie in close proximity to their associated gene/CpG, and replication of several known VNTR associations from prior targeted studies. We hypothesize that VNTRs might act to modify gene expression and epigenetics via several different mechanisms. These include modifying the structural properties of the DNA and/or chromatin fiber, changing the number of binding sites for DNA and/or chromatin-associated factors, or altering spacing between regulatory elements and their targets.

Using conditional analysis where we removed the effect of known SNV QTLs for the same gene or CpG that was associated with VNTR copy number, we show that many of the signals we detected are not simply driven by linkage disequilibrium between VNTRs and flanking SNVs. We also investigated stratification of VNTRs using diverse human populations. As selection resulting from differing environmental pressures represents a potential mechanism leading to high population divergence, elevated VST can be an indicator of possible selective effects acting on VNTR copy number. We observed multiple examples of putatively functional eVNTRs and mVNTRs that showed population-specific expansion or contraction. By annotating eVNTRs with possible human traits that they might influence based on the genes they regulate, we found that eVNTRs with elevated V_ST_ levels were enriched for putative phenotype associations. Finally, we also observed that eVNTRs and mVNTRs were enriched for tandem repeats that have undergone human-specific expansions in copy number.^48^ Overall, these data provide strong evidence to support the notion that copy number variation of some VNTR loci exerts a regulatory effect on the local genome, is likely associated with a wide variety of human traits and disease susceptibilities, and similar to single nucleotide and other types of structural variation, has likely been subject to selective pressures during recent evolutionary history.^53–55^

The majority of VNTRs we assayed using read-depth were relatively large, exceeding the read length of Illumina WGS (mean motif size 116bp, mean span of repeat tract in reference genome 353bp). Due to their size and tandemly repeated nature, copy number variation of VNTRs is difficult to assay in Illumina WGS data using standard tools for genotyping structural variants. By direct comparison with VNTR genotypes derived from long-read sequencing, we observed that *CNVnator* generally provides relatively good estimates of relative diploid VNTR copy number for the majority of VNTRs in the genome. In contrast, other published tools for genotyping VNTRs are either limited to only being able to genotype alleles that are shorter than the sequencing read length, or in our hands performed poorly for the set of VNTRs we assayed.^11,12^

However, the use of read depth does have some major limitations. First, read depth does not provide any allelic information, and only yields a relative estimate of total copy number from the sum of both alleles. For example, a heterozygous individual with alleles of two and eight repeats (total n=10) will be indistinguishable from an individual who is homozygous for an allele with five repeats (total n=10). Also the use of read depth does not differentiate between specific repeat motifs with divergent sequence that may independently vary in copy number, as has been observed to occur at some VNTRs.^4^ Furthermore, in the case that a repeat motif strongly diverges from those that are represented in the reference genome, these might be poorly measured or missed entirely, as mapping of reads to a VNTR is based on alignment to the reference sequence. Given that TR loci are frequently misassembled or collapsed during genome assembly, it is therefore likely that our study has not effectively assessed some fraction of VNTRs that are poorly represented in the current reference genome.^48,56^ Ongoing efforts to improve and diversify the human reference genome will likely provide a more complete ascertainment of VNTRs that are present in the human population.^48,57^ The use of read depth to genotype VNTRs can also potentially be confounded where a VNTR is contained within a larger underlying copy number variation, or through batch effects in WGS data. However, here we applied stringent quality control steps to remove such confounders, and through visualization of the underlying data at individual VNTRs and large-scale replication in an independent cohort, we minimized the possibility that these significantly influenced our results.

Other limitations of our analysis are that we were only able to assay CpG methylation in whole blood, and the measurements of DNA methylation that we used were based on methylation arrays, which only assay a small fraction of all CpGs in the genome. Furthermore, by using linear regression in our association analysis, we tested a model in which the relationship between expression/DNA methylation and VNTR copy number is presumed to be linear, and therefore we had limited power to identify more complex non-linear effects of different VNTR alleles that have been observed at some TR loci.^58^

Our study provides an initial map of putatively functional VNTRs, and hints that future studies of tandem repeat variation will likely yield novel insights into the genetic basis of human phenotypes that have been largely ignored in the era of SNV-based GWAS. In the future, we postulate that the use of long-read sequencing approaches that provide improved genotyping of VNTRs will lead to deeper insights into the effects of this class of structural variation on diverse human traits.

## DESCRIPTION OF SUPPLEMENTAL DATA

Supplemental Data include 9 Supplemental Figures, and 7 Supplemental Tables.

## ACKNOWLEDGEMENTS

This work was supported by NIH grant NS105781 to AJS, NIH predoctoral fellowship NS108797 to OLR, and American Heart Foundation Postdoctoral Fellowship 18POST34080396 to AMT. Research reported in this paper was supported by the Office of Research Infrastructure of the National Institutes of Health under award number S10OD018522. The content is solely the responsibility of the authors and does not necessarily represent the official views of the National Institutes of Health. This work was supported in part through the computational resources and staff expertise provided by Scientific Computing at the Icahn School of Medicine at Mount Sinai.

The Genotype-Tissue Expression (GTEx) Project was supported by the Common Fund of the Office of the Director of the National Institutes of Health (commonfund.nih.gov/GTEx). Additional funds were provided by the NCI, NHGRI, NHLBI, NIDA, NIMH, and NINDS. Donors were enrolled at Biospecimen Source Sites funded by NCI\Leidos Biomedical Research, Inc. subcontracts to the National Disease Research Interchange (10XS170), Roswell Park Cancer Institute (10XS171), and Science Care, Inc. (X10S172). The Laboratory, Data Analysis, and Coordinating Center (LDACC) was funded through a contract (HHSN268201000029C) to the The Broad Institute, Inc. Biorepository operations were funded through a Leidos Biomedical Research, Inc. subcontract to Van Andel Research Institute (10ST1035). Additional data repository and project management were provided by Leidos Biomedical Research, Inc.(HHSN261200800001E). The Brain Bank was supported supplements to University of Miami grant DA006227. Statistical Methods development grants were made to the University of Geneva (MH090941 & MH101814), the University of Chicago (MH090951, MH090937, MH101825, & MH101820), the University of North Carolina - Chapel Hill (MH090936), North Carolina State University (MH101819), Harvard University (MH090948), Stanford University (MH101782), Washington University (MH101810), and to the University of Pennsylvania (MH101822). The datasets used for the analyses described in this manuscript were obtained from dbGaP at http://www.ncbi.nlm.nih.gov/gap through dbGaP accession number phs000424.v8.p2.

The Gabriella Miller Kids First Pediatric Research Program (Kids First) was supported by the Common Fund of the Office of the Director of the National Institutes of Health (www.commonfund.nih.gov/KidsFirst). Baylor College of Medicine’s Human Genome Sequencing Center was awarded an administrative supplement (3U54HG003273-12S1) to sequence congenital cohort samples submitted by Christine Seidman through the Kids First program (1X01HL132370). The data analyzed and reported in this manuscript were accessed from dbGaP (www.ncbi.nlm.nih.gov/gap; accession number phs001138). Additional funds from the NHLBI grants U01HL098123, U01HL098147, U01HL098153, U01HL098162, U01HL098163, and U01HL098188 supported the assembly of the Pediatric Cardiac Genomics Consortium (https://benchtobassinet.com/About/AboutPCGC) cohort, and collection of the phenotypic data and samples (PMID: 23410879).

Data used in the preparation of this article were obtained from the Parkinson’s Progression Markers Initiative (PPMI) database (www.ppmiinfo.org/data). For up-to-date information on the study, visit www.ppmiinfo.org. PPMI, a public-private partnership, is funded by the Michael J. Fox Foundation for Parkinson’s Research and funding partners, a full list of which can be found at www.ppmiinfo.org/fundingpartners.

## WEB RESOURCES

1000 Genomes Project high-coverage WGS data, https://www.internationalgenome.org/data-portal/data-collection/30x-grch38

Database of Genotypes and Phenotypes (dbGaP), https://www.ncbi.nlm.nih.gov/gap/

Gene Expression Omnibus (GEO), https://www.ncbi.nlm.nih.gov/geo/

GWAS catalog, https://www.ebi.ac.uk/gwas/

OMIM, http://www.omim.org/

Parkinson’s Progression Markers Initiative (PPMI), https://www.ppmi-info.org/

UCSC Genome Browser, http://genome.ucsc.edu

## Supplementary Tables

**Table S1.** 404 samples from the GTEx cohort that were used for association analysis of VNTR copy number with gene expression.

**Table S2.** 14 individuals with both Illumina and PacBio whole genome sequencing data used to assess the performance of VNTR genotyping with *CNVnator* and *adVNTR*.

**Table S3.** Comparison of VNTR genotypes derived from PacBio WGS compared with those from *CNVnator* and *adVNTR* derived from Illumina WGS in 14 samples.

**Table S4.** Results of association analysis of VNTRs with gene expression in 48 tissues from the GTEx discovery cohort. Additional annotation includes putative trait associations of the associated gene from the GWAS catalog, *PrediXcan*, and OMIM, conditional analysis based on the lead eQTL SNV, population stratification, and human specific repeat expansions.

**Table S5.** Results of association analysis of VNTRs with CpG methylation in the PCGC discovery cohort. Additional annotation includes conditional analysis based on the lead mQTL SNV, population stratification, and human specific repeat expansions.

**Table S6.** Results of association analysis of VNTRs with gene expression in the PPMI replication cohort.

**Table S7.** Results of association analysis of VNTRs with CpG methylation in the PPMI replication cohort.

