## Supplemental Figures for "Pervasive *cis* effects of variation in copy number of large tandem repeats on local epigenetics and gene expression"

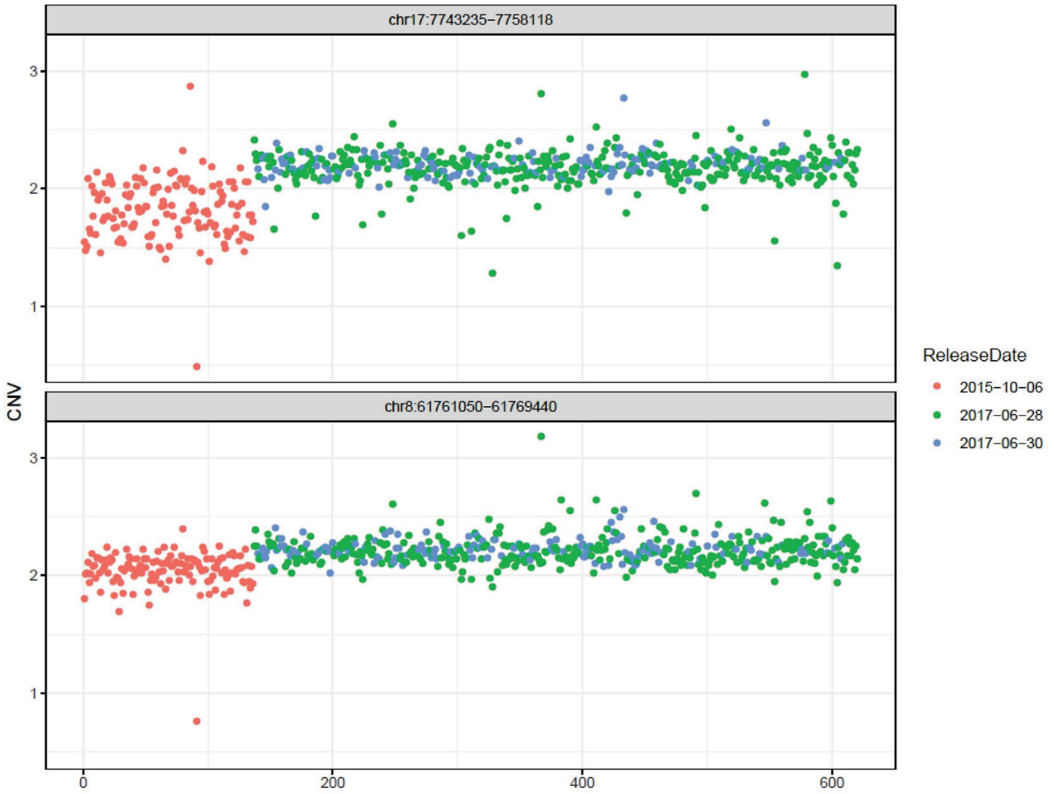


**Figure S1. Batch effects in *CNVnator* copy number estimates from WGS in the GTEx cohort.** Using *CNVnator* data for two highly constrained genes that should remain copy-number invariant in the normal population (*KDM6B* [MIM: 611577], chr17:7,839,904-7,854,796 and *CHD7* [MIM: 608892], chr8:60,678,740-60,868,028, hg38), we observed a strong batch effect in the GTEx cohort, whereby copy numbers derived from WGS data with release date October 6$^{th}$ 2015 (*red points*) were systematically shifted compared to later data releases (*blue and green points*). Based on these observations, we removed from further analysis all 135 samples with release October 6th 2015.

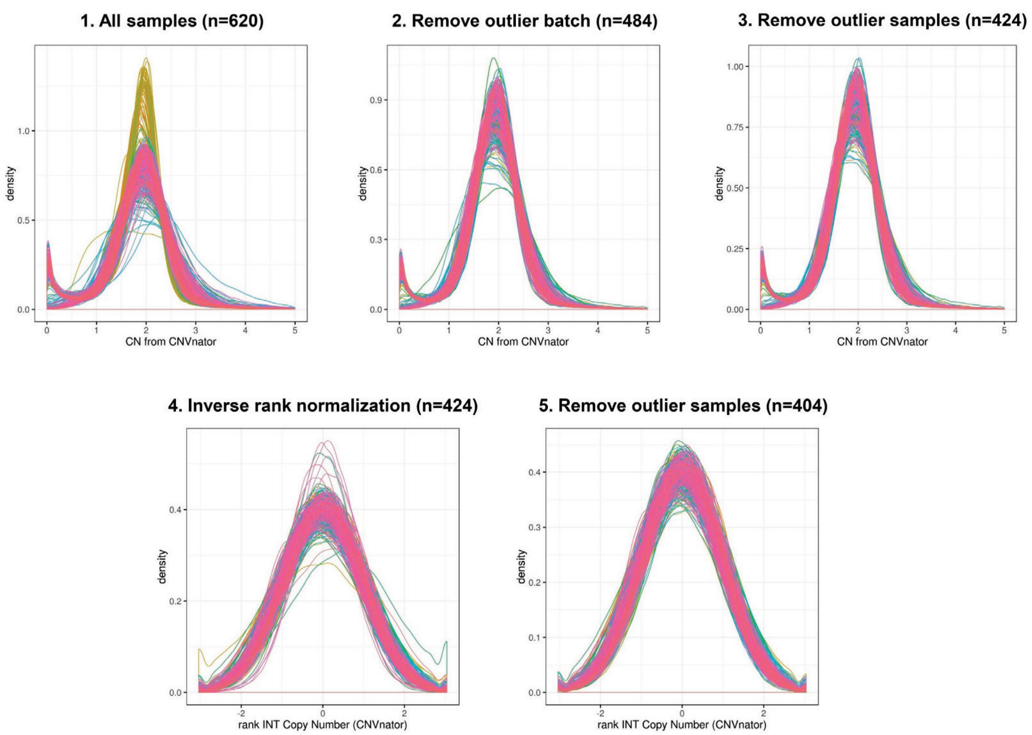


**Figure S2. VNTR copy number distributions within the GTEx cohort after sample filtering and data normalization.** Based on *CNVnator* copy number estimates of 89,893 VNTRs, we generated density plots at each step of quality control and normalization. We initially analyzed WGS data from 620 individuals, but after removal of batch effects, samples that were consistent outliers at invariant constrained genomic loci, or outliers for VNTR copy number by principal component analysis and density plots, we used a final cohort of 404 samples in our analysis.

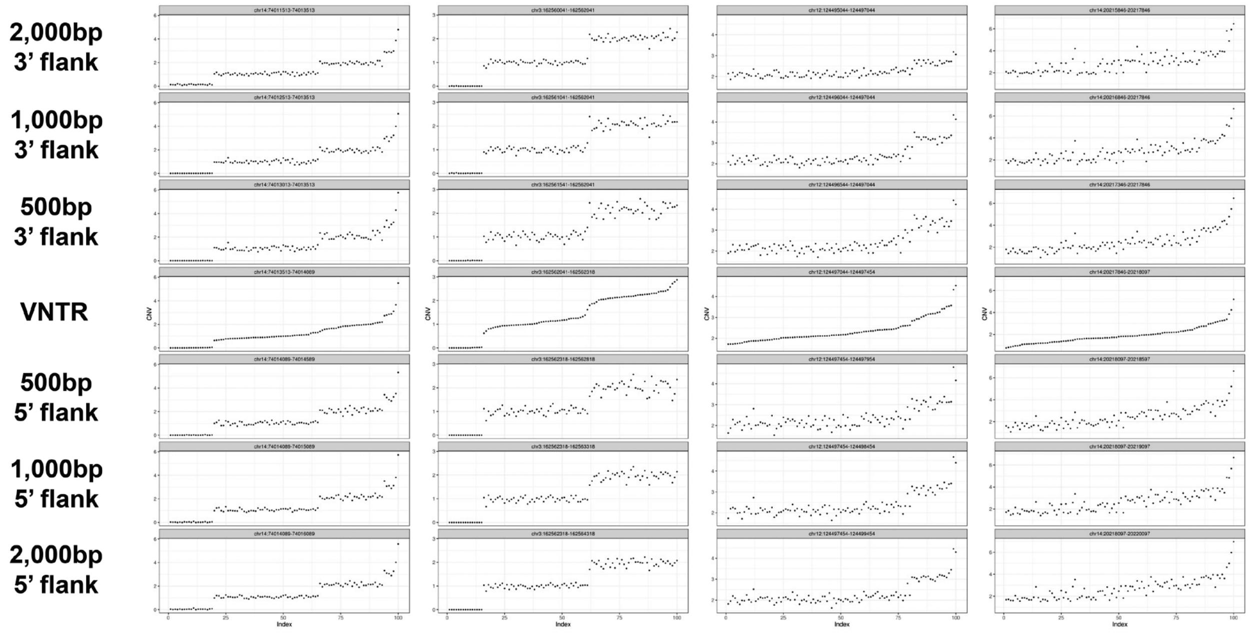


**Figure S3. VNTR copy number estimates using *CNVnator* can be confounded by the presence of larger underlying CNVs.** In situations where a VNTR is embedded within a larger copy number variable region, copy number estimates for the VNTR based on *CNVnator* read depth can be influenced by underlying variations of the wider region. To identify VNTRs that were subject to this confounder, we analyzed the 3' and 5' 500bp, 1kb and 2kb regions flanking each VNTR using *CNVnator*, and then correlated the values of the 1kb flanks with VNTR copy number. Shown are data from four representative loci that were removed from further analysis. Within each locus, samples are ordered based on the estimated VNTR copy number, revealing that the observed estimates of VNTR copy number are highly correlated with variation in the flanking regions, and likely simply reflect a larger underlying CNV, rather than changes in length of the VNTR array itself.

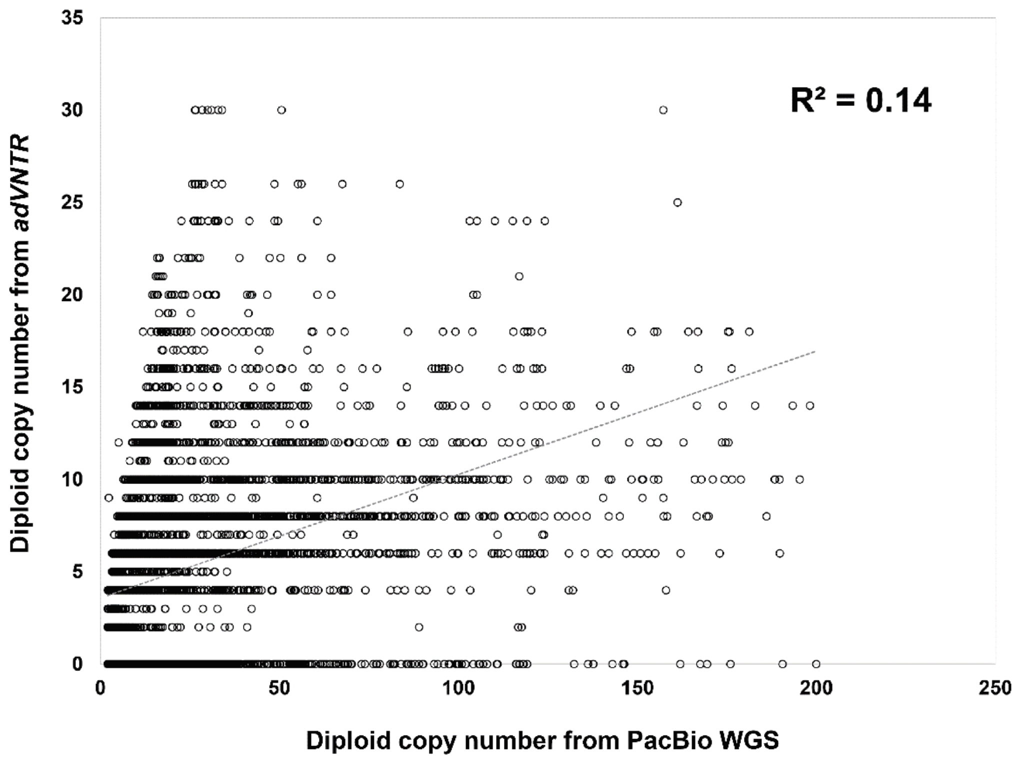


**Figure S4. Poor performance of *adVNTR* for genotyping VNTRs.** Using data for the same set of 1,891 VNTRs in 14 individuals as shown in Figure 1B (see Methods), we used *adVNTR* to generate VNTR genotypes. When compared with direct genotypes generated from PacBio long-read WGS in these same individuals, we observed an $R^2$=0.14, indicating generally poor accuracy of this tool when applied to this set of VNTRs (Table S3).

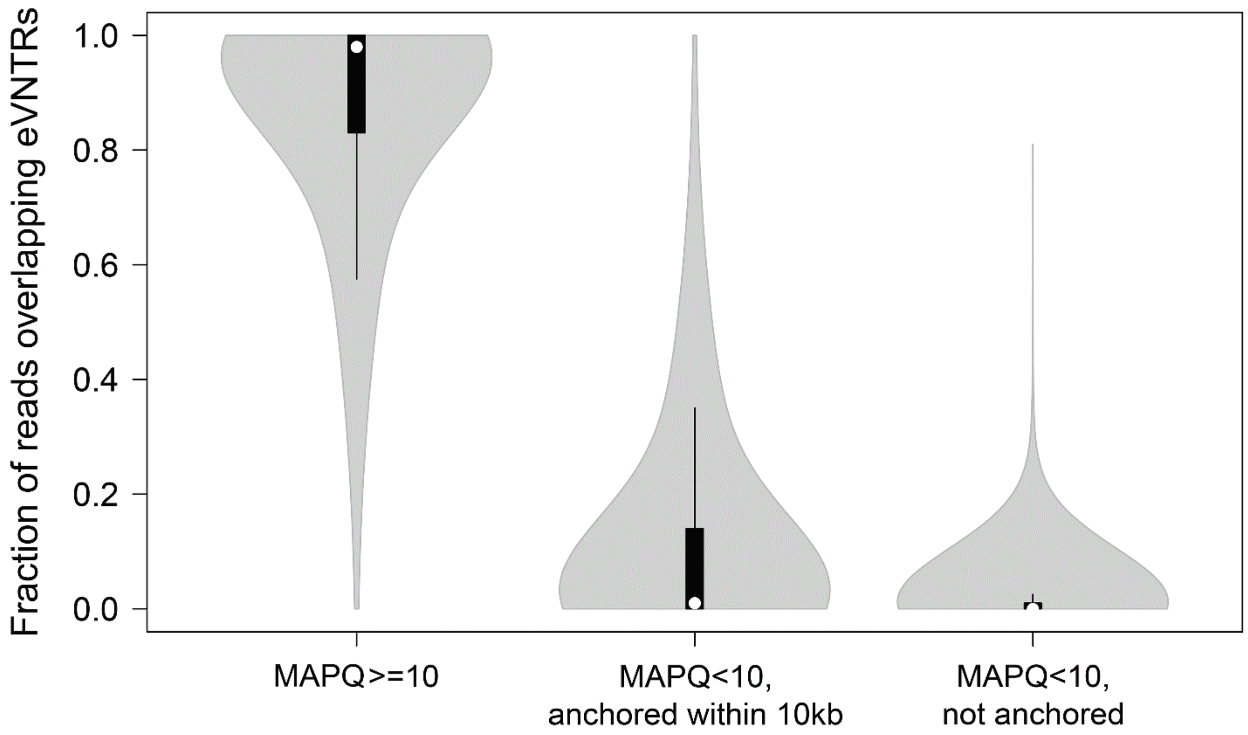


**Figure S5. Assessment of the reliability of Illumina reads mapping to VNTR loci.** We analyzed Illumina reads mapping to 2,980 GTEx eVNTRs in a Yoruban sample (NA18874), classifying them into three categories: (i) MAPQ≥10, (ii) MAPQ<10, but with a mate-pair that mapped reliably within ±10kb. (iii) MAPQ<10, without a mate pair that was anchored within ±10kb. Violin plots show the fraction of reads in each of these three categories at each eVNTR locus. Overall, copy number estimates for the vast majority of eVNTRs were based on reliably mapped reads, with only a single eVNTR containing >50% of unreliably mapped reads. Within each violin, the median is indicated by the white dot, box limits indicate the 25th and 75th percentiles, and whiskers extend 1.5 times the interquartile range from the 25th and 75th percentiles.

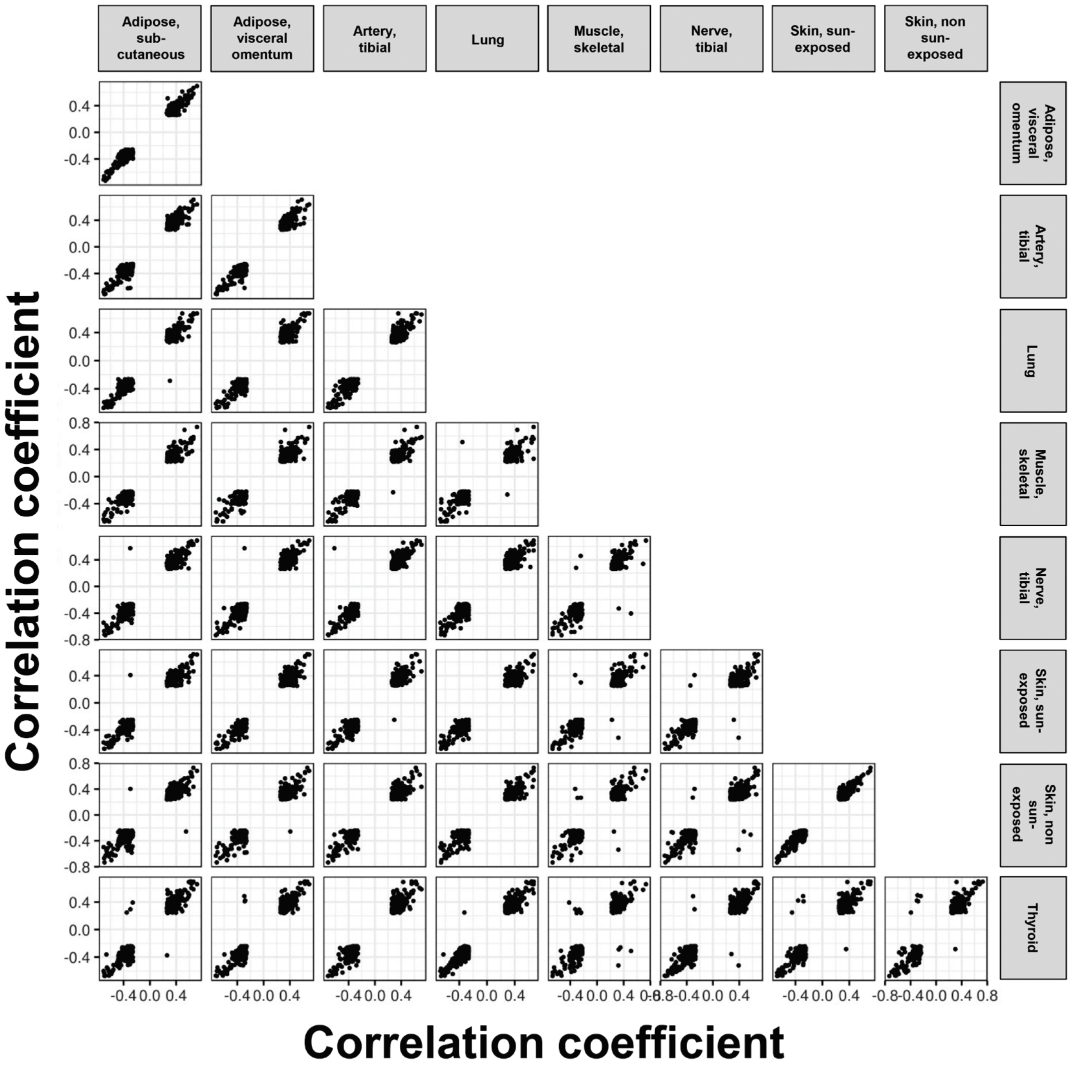


**Figure S6. Pairwise correlation patterns of significant eVNTR:gene associations across eight tissues.** Each point shows the R values of an individual gene:eVNTR pair that was significant in both tissues. In nearly all cases, the directionality of the observed associations are concordant among different tissues, with only 0.6% of eVNTR:gene pairs showing opposite direction of effect in different tissues, consistent with our results representing genuine associations.

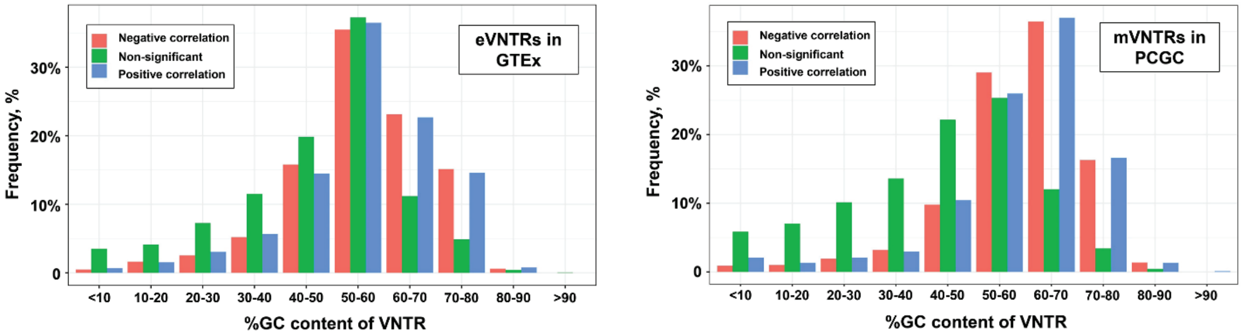


**Figure S7. Significant eVNTRs and mVNTRs are biased towards higher GC content.** We observed that both putatively functional eVNTRs and mVNTRs showed a clear trend to be composed of motifs with higher GC-content than the background of all VNTRs tested. This trend was stronger for mVNTRs, which is consistent with the Illumina 850k array preferentially sampling CpG in GC-rich regions of the genome, and the observation that most mVNTRs are located physically close (<5kb) from the CpGs that they associated with.

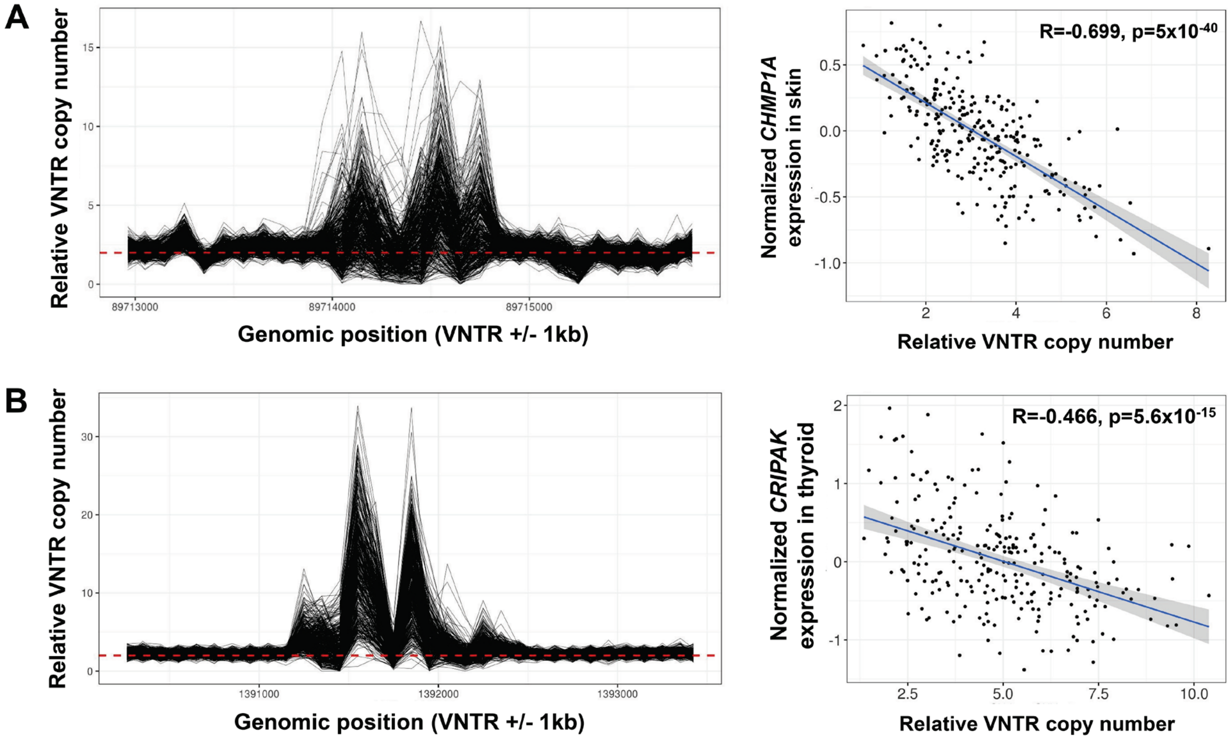


**Figure S8. Two example eVNTR loci detected in the GTEx cohort. (A)** A 42mer repeat (chr16:89,647,518-89,648,445, hg38) located intronic within *CHMP1A* [MIM: 164010] associates negatively with *CHMP1A* expression in multiple tissues (shown is data from skin, sun exposed lower leg). **(B)** A 45mer repeat (chr4:1,397,437-1,398,660) located 1.4 kb downstream of *CRIPAK* [MIM: 610203] associates negatively with *CRIPAK* expression in multiple tissues (shown is expression data from thyroid). *CNVnator* locus plots show estimated copy number per 100bp bin over the VNTR region, extending 1kb each side, with the red dashed line indicating diploid copy number equal to that of the reference genome.

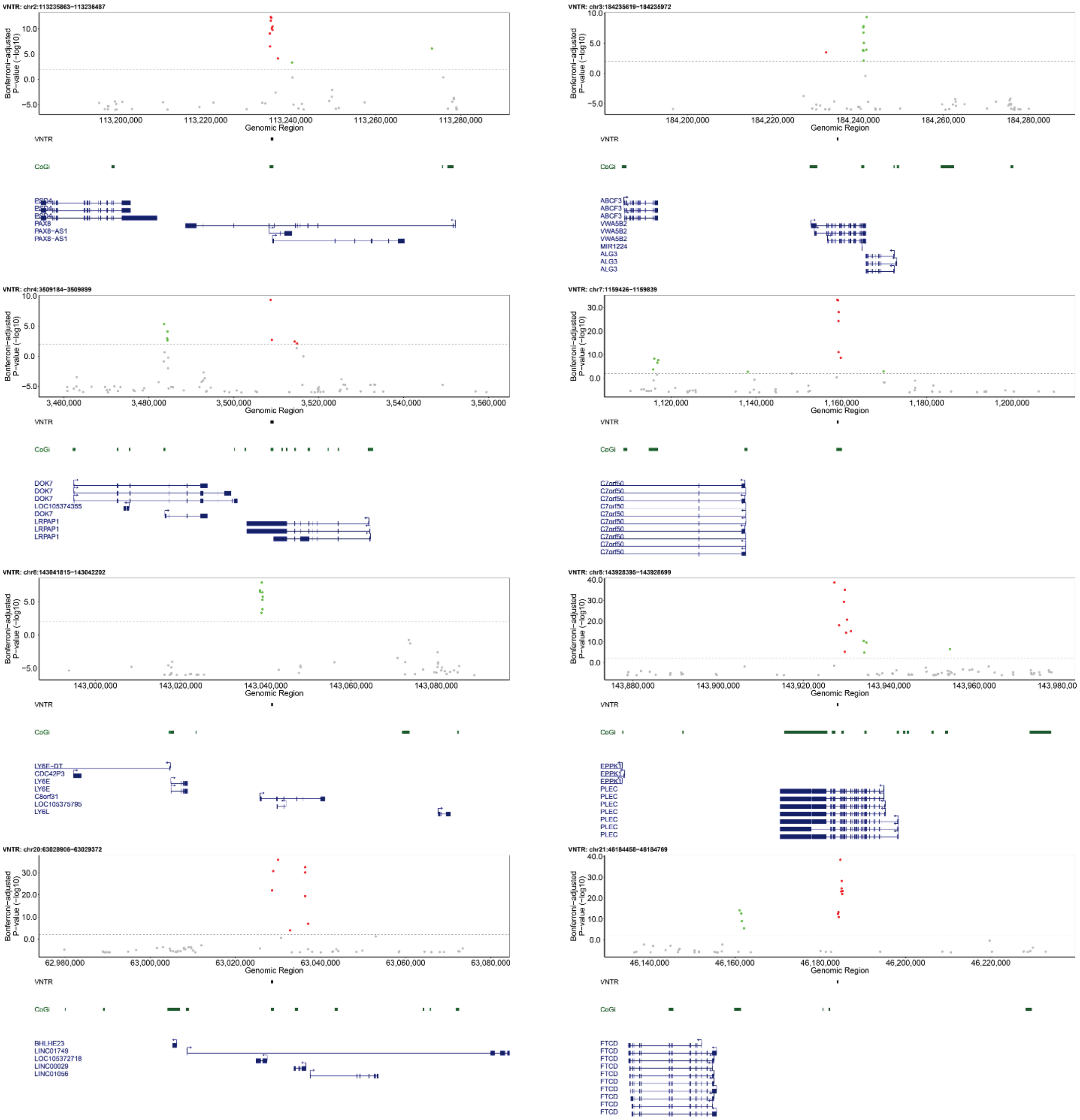


**Figure S9. Eight example mVNTR loci detected in the PCGC cohort.** Each plot shows associations between copy number of a VNTR and CpG methylation within ±50kb (hg38 coordinates). The horizontal dashed line indicates the significance threshold (p<0.01 after Bonferroni correction for the number of pairwise tests performed genome-wide), with significant CpGs shown in color, with red representing positive correlations with VNTR copy number, and green indicating negative correlations. The location of the VNTR is indicated by the horizontal black bar in the center below each plot. Underneath each plot are shown the location of CpG islands (green bars) and Refseq genes (blue).
